# cAMP-independent control of twitching motility in *Pseudomonas aeruginosa*

**DOI:** 10.1101/116954

**Authors:** Ryan N.C. Buensuceso, Martin Daniel-Ivad, Sara L.N. Kilmury, Hanjeong Harvey, P. Lynne Howell, Lori L. Burrows

## Abstract

FimV is a *Pseudomonas aeruginosa* inner membrane hub protein that modulates levels of the second messenger, cyclic AMP (cAMP), through activation of the adenylate cyclase, CyaB. Although type IVa pilus (T4aP)-dependent twitching motility is modulated by cAMP levels, mutants lacking FimV are twitching impaired, even when exogenous cAMP is provided. Here we further define FimV’s cAMP-dependent and -independent regulation of twitching. We confirmed that the response regulator of the T4aP-associated Chp chemotaxis system, PilG, required both FimV and the CyaB regulator, FimL, to activate CyaB. However, in cAMP-replete backgrounds - lacking the cAMP phosphodiesterase CpdA or the CheY-like protein PilH, or expressing constitutively-active CyaB - *pilG* and *fimV* mutants failed to twitch. Both cytoplasmic and periplasmic domains of FimV were important for its cAMP-dependent and -independent roles, while its septal peptidoglycan-targeting LysM motif was required only for twitching motility. Polar localization of the sensor kinase PilS, a key regulator of transcription of the major pilin, was FimV-dependent. However, unlike its homologues in other species that localize flagellar system components, FimV was not required for swimming motility. These data provide further evidence to support FimV’s role as a key hub protein that coordinates the polar localization and function of multiple structural and regulatory proteins involved in *P. aeruginosa* twitching motility.

**IMPORTANCE:** *Pseudomonas aeruginosa* is a serious opportunistic pathogen. Type IVa pili (T4aP) are important for its virulence, because they mediate dissemination and invasion via twitching motility, and are involved in surface sensing which modulates pathogenicity via changes in cAMP levels. Here we show that the hub protein FimV and the response regulator of the Chp system, PilG, regulate twitching independently of their roles in modulation of cAMP synthesis. These functions do not require the putative scaffold protein FimL, proposed to link PilG with FimV. PilG may regulate asymmetric functioning of the T4aP system to allow for directional movement, while FimV appears to localize both structural and regulatory elements – including the PilSR two-component system – to cell poles for optimal function.

## INTRODUCTION

Type IV pili (T4P) are polar filamentous surface appendages made by a broad range of bacteria and archaea (1, 2). They can be divided into two sub-families, type IVa (T4aP) and type IVb (T4bP), which – though clearly related – differ in their pilin subunits and assembly system architectures (2). T4aP are involved in several processes including DNA uptake, surface attachment, and twitching motility (3-5). During twitching motility, T4aP undergo repeated cycles of assembly and disassembly, acting as molecular grappling hooks to pull the cells along surfaces. Well-studied T4aP model species include *Neisseria spp*., *Myxococcus xanthus*, and *Pseudomonas aeruginosa* (6, 7). Although core structural components of the T4aP assembly machinery and pilus fibre are shared, each species has unique regulatory elements that control the function of the T4aP machinery in response to their specific environmental requirements. Without these regulatory proteins, the bacteria make non-functional T4aP systems (8-11).

The *P. aeruginosa* Chp system is a putative chemosensory system that controls both twitching motility and intracellular levels of the second messenger, cyclic adenosine monophosphate (cAMP) (12-14). It resembles the well-studied Che system of *E. coli*, but lacks a CheZ-like phosphatase. Rather, similar to *Sinorhizobium meliloti* (15), it has two CheY-like response regulators, PilG and PilH (14, 16). PilG is proposed to regulate activation of CyaB and pilus extension (17), while PilH has been proposed to be either a phosphate sink that limits downstream signalling through PilG in lieu of a phosphatase (17, 18), or a separate response regulator controlling function of the T4aP retraction ATPase, PilT (12).

The Chp system positively regulates intracellular levels of cAMP by activating the major adenylate cyclase, CyaB (17). Deletion *of pilG* results in decreased cAMP, surface piliation, and twitching motility, while *pilH* mutants have increased cAMP and surface piliation but decreased twitching relative to wild type (17). Supplementation of a *pilG* mutant with exogenous cAMP restored surface piliation but not twitching motility (17), suggesting that PilG regulates pilus biogenesis and function by at least two pathways. A recent study (18) showed that of the two proteins, PilH is the preferred target of ChpA phosphorylation, consistent with its proposed role as a phosphate sink. Decreased twitching motility in the *pilH* background may reflect hyperphosphorylation of PilG, perturbation of the chemotactic response, and uncoordinated movement.

Important for *P. aeruginosa* virulence is its ability to switch from a planktonic to sessile state when cells contact surfaces (19, 20). T4aP-mediated surface interaction is proposed to lead to signalling through the Chp system, upregulating surface-associated virulence phenotypes by increasing intracellular levels of cAMP (20). Vfr (virulence factor regulator) binds cAMP and modulates the expression of >200 genes, including the type II secretion system (T2SS) and its effectors, and T4aP assembly components including the motor ATPases PilBTU, the alignment subcomplex PilMNOP, the secretin PilQ, and the PilSR two-component system that regulates PilA levels (21). This regulatory circuitry allows for just-in-time expression of components required for a surface-associated lifestyle in response to surface contact.

FimV is also required for T4aP function and CyaB activation (17), and is proposed to link into the Chp system via the cytoplasmic protein, FimL (22). FimV is a 97 kDa inner-membrane protein with one transmembrane segment. Its periplasmic domain contains a lysin (LysM) motif that binds peptidoglycan (PG) (23), and its cytoplasmic domain contains three discontinuous tetratricopeptide repeat (TPR) motifs involved in protein-protein interactions (24). FimV homologs have been identified in other T4P-producing bacteria (10) although their overall sequence identity is low, with the most conserved features being the LysM motif (COG3170), the single transmembrane segment, and a highly conserved cytoplasmic “FimV C-terminal domain” – TIGR03504 – encompassing a single TPR repeat and capping helix (25).

FimV homologs have been characterized in several species (8-11, 26–28), but their functions are not necessarily conserved. Deletion of FimV in *Legionella pneumophila* resulted in loss of twitching motility and cell elongation, while deletion of the *Neisseria meningitidis* FimV homolog TspA led to decreased host cell adhesion but no effect on twitching motility or surface piliation. The *Vibrio cholerae* homolog, HubP, functions as a protein interaction hub, although its role is not limited to T4P localization. Deletion of HubP altered the cellular distribution of the chemotactic and flagellar machinery, and the chromosomal origin, *oriCI* (28). HubP from *Shewenella putrifaciens* is responsible for localization of the chemotactic machinery, but not the flagellar system (27). Yamaichi et al. (28) showed that polar localization of *V. cholerae* HubP was dependent on the conserved LysM motif (25). Wehbi et al. (29) showed that *P. aeruginosa fimV* mutants have decreased levels of the T4aP alignment subcomplex proteins, PilMNOP, while inframe deletion of FimV’s LysM motif resulted in fewer PilQ multimers, suggesting that PG binding is important for optimal secretin formation. A recent study (30) confirmed that FimV participates in localization of PilMNOPQ to sites of future cell division, ultimately placing T4aP assembly systems at both poles of newly divided cells.

T4aP-mediated twitching motility requires both cAMP-dependent and independent inputs (17). For example, provision of exogenous cAMP to mutants lacking PilG restored piliation but not motility, and a mutant expressing a constitutively active form of CyaB but lacking FimV failed to twitch (25). FimL was proposed to be a scaffold protein linking PilG to the C-terminal TPR motif of FimV, leading to CyaB activation, and FimV localized both FimL and PilG to cell poles (22, 25). However, of these three proteins, only FimL is dispensable for twitching motility in cAMP-replete conditions. Thus, the FimV-FimL-PilG model fails to explain the cAMP-independent roles of FimV and PilG in twitching.

Here we provide evidence supporting different cAMP-independent roles for FimV and PilG in regulation of twitching motility. We show that in addition to polar localization of FimL, PilG, and PilMNOPQ (22, 30), FimV is responsible for polar localization of PilS, the membrane-bound sensor kinase that controls *pilA* transcription. These data show that FimV plays a central role in control of twitching motility that overlaps with – but is distinct from – that of the Chp system.

## RESULTS

### FimV is required for Chp activation of CyaB

FimL, FimV, and PilG are all required for activation of CyaB, with FimL proposed to link FimV to the Chp system through PilG (17, 22, 31, 32). However, while phenotypes associated with *fimL* deletion could be rescued by deletion of *cpdA* or by increasing intracellular cAMP levels in other ways (22, 32, 33), provision of exogenous cAMP failed to restore motility in a *pilG* mutant (17). We investigated whether the cAMP-independent function of PilG also required FimV by comparing PilU levels – a proxy for intracellular cAMP levels (21, 25) – twitching, and piliation in *fimL*, *fimV*, and *pilG* single mutants or in double mutants also lacking *cpdA* to prevent degradation of endogenous cAMP (32) (Figure 1). To confirm that FimV and FimL were epistatic to PilG, we also examined *pilH fimL* and *pilH fimV* double mutants. In the absence of PilH, cells are predicted to have hyper-phosphorylated PilG, consistent with the high levels of cAMP observed in a *pilH* background (17, 18).

**Figure 1.**
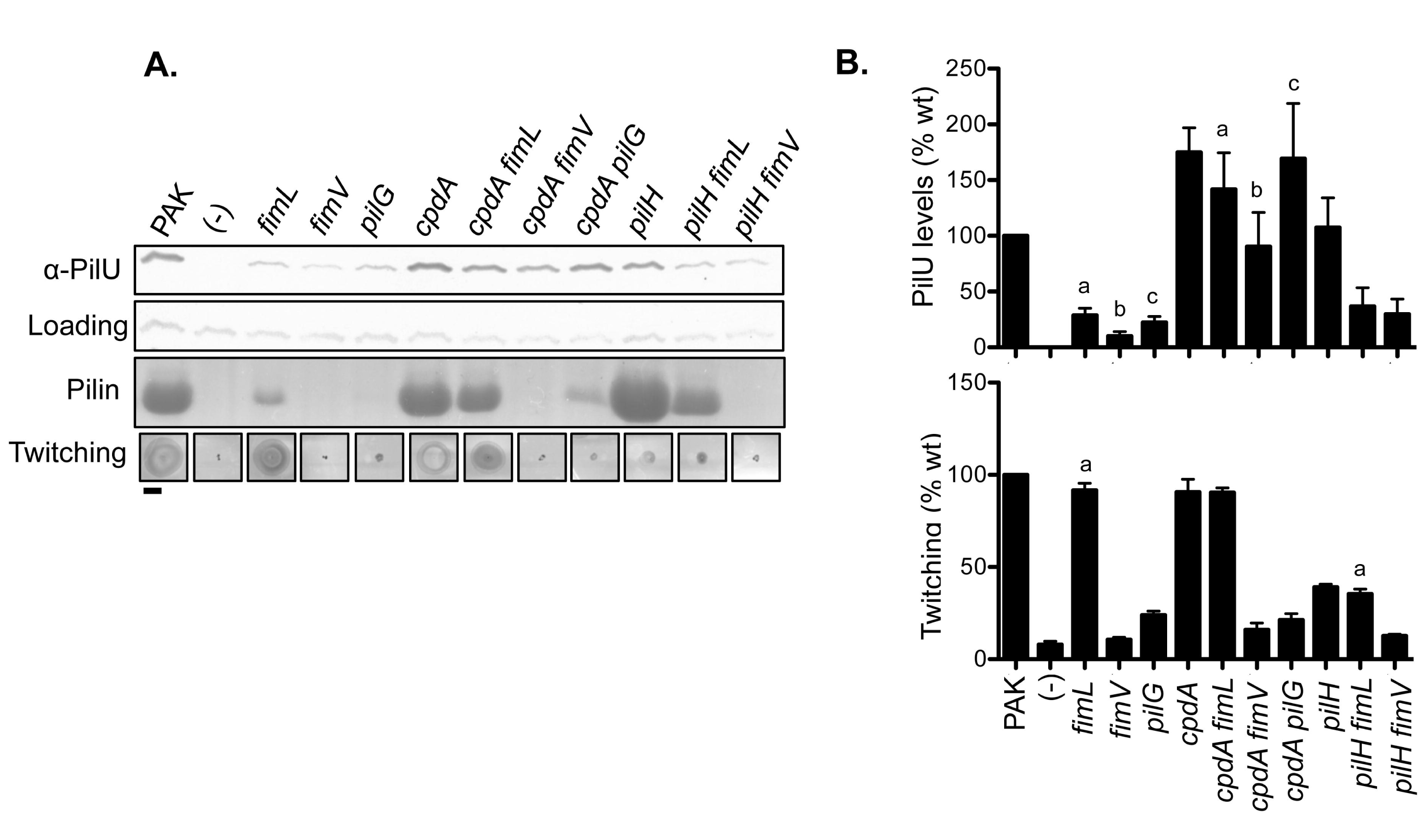
FimV and PilG have separate cAMP-independent roles in T4aP function. **A.** Loss of *fimL* reduces the levels of PilU (a proxy for intracellular cAMP levels), surface piliation, and twitching motility, and all phenotypes are restored to wild type levels in a *cdpA fimL* double mutant. In contrast, *fimV* and *pilG* mutants lack surface piliation and motility even in a *cpdA* background, while PilU levels are restored to wild type or greater. Scale bar = 1 cm. **B.** Quantification of relative PilU levels and twitching zone diameters compared to wild type PAK set to 100%; measurements are the average of n=3. Matching letters indicate statistically significant differences at p<0.05.

PilU levels were decreased in *fimL*, *fimV*, and *pilG*, to 28%, 10% and 22% of wild type, respectively, consistent with roles in regulating cAMP synthesis (10, 12, 17, 32), (Figure 1). Both *pilG* and *fimV* were twitching deficient, whil*e fimL* twitching resembled that of wild type, as reported previously (32, 34). The *cpdA* mutant had high levels of PilU, surface piliation, and wild type twitching, consistent with high cAMP levels (17, 32). Deletion of *cpdA* in *fimL*, *fimV*, or *pilG* increased PilU levels relative to the corresponding single mutants, to at least wild-type levels (Figure 1), showing that CyaB retains residual activity in those backgrounds. However, only the *cpdA fimL* mutant was motile, confirming that both PilG and FimV have cAMP-independent roles in T4aP function. Also consistent with previous reports (17), a *pilH* mutant assembled surface pili but was twitching impaired (~39% of wild type). The *pilH fimV* and *pilH fimL* double mutants had PilU levels similar to those of *fimV* and *fimL* single mutants, suggesting that despite its hyper-phosphorylation in the absence of *pilH* (18), PilG was unable to activate CyaB without FimV or FimL, confirming that all three are required for the Chp system to stimulate cAMP synthesis.

### Decreased levels of PilMNOPQ in *fimV* are due to decreased cAMP

Wehbi et al. (29) showed previously that *fimV* mutants had reduced levels of PilMNOP, and *that a fimV*_∆*LysM*_ mutant with an in-frame deletion of the LysM motif had fewer PilQ secretins. However, transcription of the *pilMNOPQ* operon is Vfr-and thus cAMP-dependent (21). To determine if any of these phenotypes were independent of cAMP, we examined levels of PilMNOPQ and PilU, and twitching motility in *fimV*, *fimV*_∆*LysM*_, a mutant encoding only the cytoplasmic domain of FimV (*fimV*_1194_), and in a *fimV cyaB-R456L* double mutant that expresses constitutively active CyaB (25, 35).

The *fimV* mutant had low levels of PilU (~23% of wild type), reflecting low cAMP levels, while the *fimV cyaB-R456L* double mutant had wild type levels of PilU (Figure 2). The *fimV*_1194_ mutant had ~29% of wild-type PilU, suggesting that even though its protein partners PilG and FimL are cytoplasmic, expression of FimV’s cytoplasmic domain alone was insufficient to activate CyaB. Complementation of *fimV* and *fimV_1194_ in trans* with a construct expressing full-length FimV increased PilU levels to ~58% and ~65% of wild type, respectively. Surprisingly, the *fimV_LysM_* mutant had ~89% of wild type PilU, suggesting that the LysM motif and thus PG binding was dispensable for CyaB activation.

**Figure 2.**
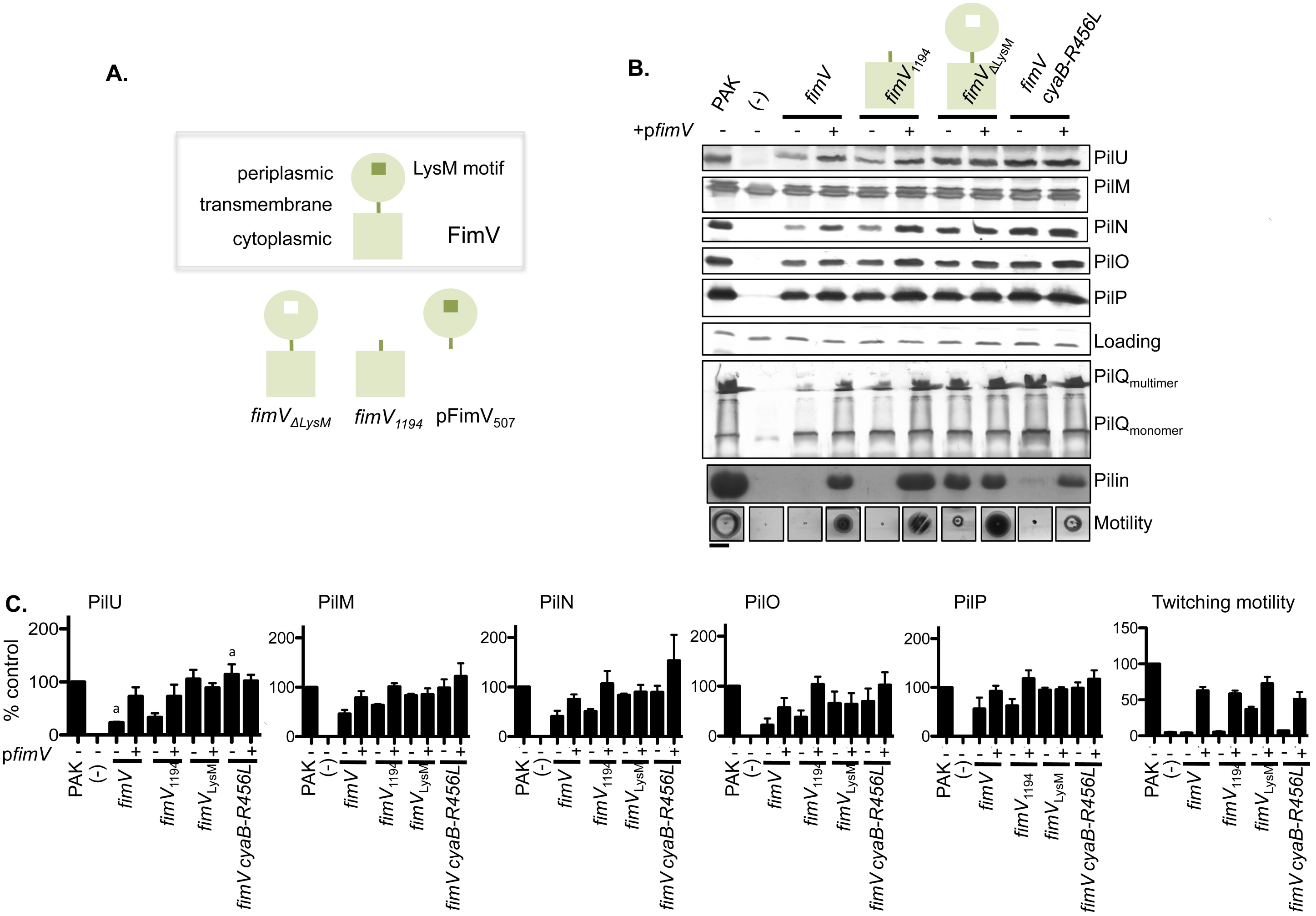
FimV, and its ability to bind PG, are essential for motility. **A.** Schematic of FimV fragments used in this work. FimV’s periplasmic and cytoplasmic domains are connected by a single transmembrane segement. The N-terminal periplasmic domain contains a PG-binding LysM motif (green square), which is deleted in-frame in the *fimV_∆LysM_* mutant (white square). **B.** The levels of PilU (a proxy for intracellular cAMP levels; (25)), and assembly machinery components PilMNOPQ were assessed by Western blot in various *fimV* backgrounds, complemented with empty vector or with full-length *fimV;* a representative blot is shown. Surface piliation and twitching motility are shown in the bottom two rows. Scale bar = 1 cm. **C.** Quantification of protein levels and motility, n = 3. Note that PilQ multimer levels cannot be accurately quantified by this method. In the absence of *fimV*, levels of PilU, PilN, and PilO and PilQ multimers were reduced, and surface piliation and motility were lost. These phenotypes were restored by complementation with *fimV* in *trans. fimV_1194_* expressing only the cytoplasmic domain phenocopied the *fimV* deletion mutant. When FimV’s LysM motif was deleted, levels of the proteins of interest and piliation were similar to wild type, but motility remained severely impaired. CyaB requires FimV for its activity, but in the absence *of fimV* can be constitutively activated by the R456L mutation (35). However, the cells have very few surface pili and are non-motile.

The *fimV* mutant had decreased levels of PilMNOP, and few detectable PilQ multimers, and all were restored to wild type with full-length FimV. Supporting the hypothesis that their levels were dependent on Vfr and cAMP, the *fimV cyaB-R456L* double mutant had wild type levels of PilMNOPQ. Despite this, the *fimV cyaB-R456L* double mutant had no recoverable surface pili (Figure 2) and could not twitch, confirming a cAMP-independent role(s) for FimV in pilus assembly and twitching motility. *The fimV_1194_* mutant had low levels of PilMNOP and no detectable PilQ multimers, but these could be rescued by complementation with full length FimV. fimV_∆LysM_ had essentially wild type PilMNOPQ levels, and could assemble surface pili (Figure 2); however, twitching was ~37% of wild type. The motility defect in *fimV*_∆LysM_ suggests that PG binding is important for FimV’s cAMP-independent function(s).

Wehbi et al. (29) showed that the *fimV_1194_* mutant (which expresses the cytoplasmic domain of FimV) was unable to twitch, but motility could be rescued by complementation with a plasmid expressing only the periplasmic domain of FimV (residues 1-507, pFimV_507_), suggesting that together, the two FimV fragments could restore function without being physically connected. *A fimV* deletion mutant complemented with empty vector or pFimV_507_ had similar PilU levels, suggesting that the periplasmic domain alone is not sufficient to activate CyaB (Figure 3A). Unexpectedly, despite its ability to restore motility in the *fimV_1194_* background (Figure 3B), pFimV_507_ did not significantly increase PilU levels in that background, suggesting that the cytoplasmic and periplasmic domains cannot activate CyaB efficiently when they are not covalently linked. However, CyaB activation is not strictly required for twitching, as both *fimL* (Figure 1) and *cyaAB* mutants have low cAMP levels and piliation, but near wild-type motility (17, 32). We next tested if FimV’s periplasmic domain played a cAMP-independent role in twitching by complementing the *fimV cyaB-R456L* mutant with pFimV_507_. pFimV_507_ failed to restore twitching in *fimV cyaB-R456L* (Figure 3B), suggesting that the cytoplasmic region of FimV plays a cAMP-independent role in motility. Taken together, the data show that the cAMP-independent role(s) of FimV requires both domains.

**Figure 3.**
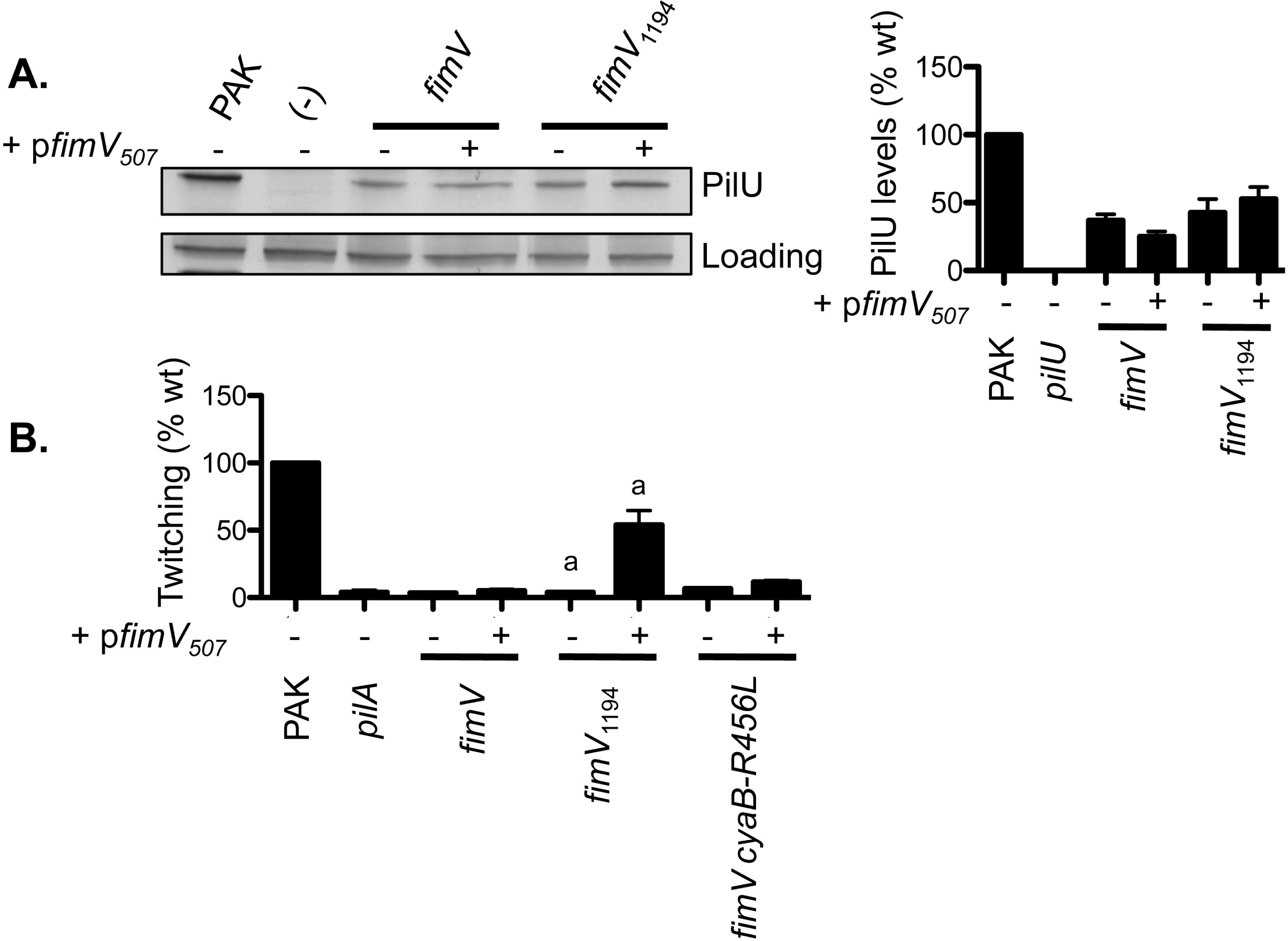
CyaB activation - but not twitching motility - requires that FimV’s cytoplasmic and periplasmic domains be connected. **A.**Representative Western blot of whole cell lysates with anti-PilU antiserum, and quantification of PilU levels (a intreacellular for cAMP levels) by densitometry, n = 3. The reduced levels of PilU in *fimV* and *fimV_1194_* mutants are not restored to wild type by *in trans* expression of the N-terminal domain (FimV_507_). **B.** Quantification of twitching motility. As reported previously (29), expression of pFimV_50_7 *in trans* complements motility in the *fimV1194* mutant that expresses the cytoplasmic domain of FimV. It cannot complement motility of a *fimV* mutant, even when cAMP levels are restored in a background expressing constitutively-active CyaB-R456L.

### FimV is required for PilS localization

The *V. cholerae* homolog of FimV, HubP, interacts with multiple proteins and has broad regulatory function (28). FimV is required for bipolar localization of PilG and FimL and the T4aP structural proteins PilMNOPQ, but not the Chp methyl-accepting chemotaxis protein (MCP) PilJ (22, 30). To determine if FimV was required for localization of other T4aP regulators, we examined its effects on localization of PilS (36, 37), the histidine sensor kinase component of the PilRS two-component system that regulates *pilA* transcription in response to changes in PilA levels in the inner membrane (38).

In wild-type cells, PilS-YFP was localized to both poles (Figure 4) as reported previously (36), while in the absence of FimV, PilS-YFP was diffuse in the inner membrane. Interestingly, the localization pattern of PilS-YFP in *fimV*_∆LysM_ was similar to wild type, suggesting that PG-binding was not critical for PilS localization. However, PilS-YFP was mislocalized in *fimV_1194,_* suggesting that the cytoplasmic domain is insufficient for PilS localization. Finally, because PilG also has cAMP-independent effects on motility (17) (Figure 1), we examined PilS-YFP localization in a *pilG* mutant. Localization was similar to wild type, suggesting that PilG and FimV have distinct cAMP-independent roles in motility (Figure 4). These data also suggest that PilS localization is independent of intracellular cAMP concentration.

**Figure 4.**
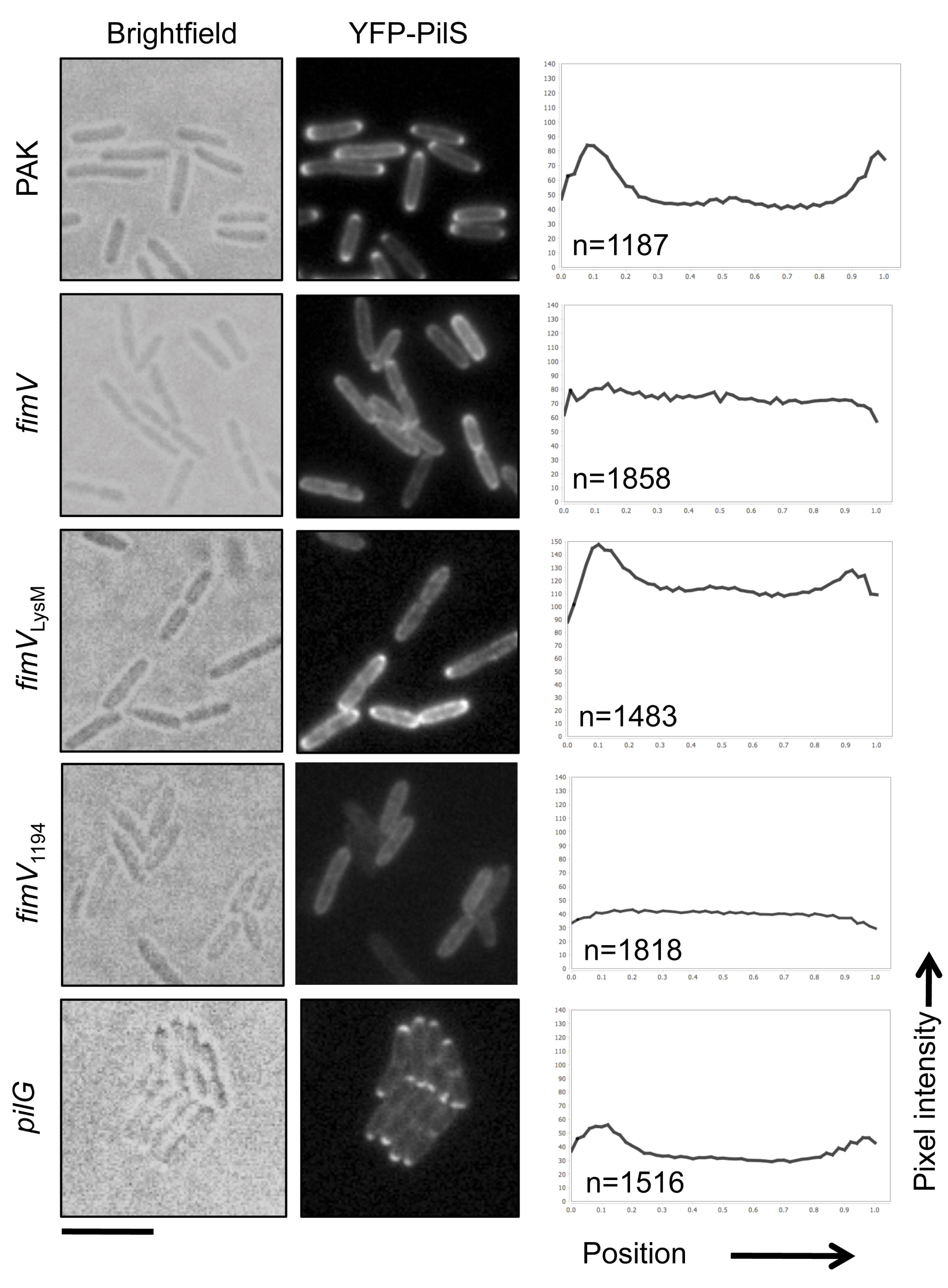
PilS localization is dependent on FimV. Brightfield and fluorescence microscopy was used to image PilS-YFP localization in the wild type, *fimV* mutants, or a *pilG* mutant and the average pixel intensity of the YFP signal along the long axis of the cell was quantified using MicrobeJ (48). The number of cells averaged for each population is shown on the graph. PilS-YFP is localized to the poles in wild type cells but fluorescence becomes circumferential in the *fimV* and *fimV_1194_* backgrounds. In a mutant expressing FimV with an in-frame deletion of its LysM peptidoglycan-binding motif, fluorescence is polar but patchy circumferential fluorescence is also visible. PilS-YFP localization remains polar in a *pilG* mutant. Scale bar = 5 μm.

### FimV deletion does not affect swimming motility

As the *V. cholerae* and *S. putrefaciens* homologs of FimV modulate swimming motility (27, 28), we tested whether loss of *fimV* impaired swimming in *P. aeruginosa.* We saw no effect of FimV deletion on swimming motility (Figure 5), suggesting that it is not essential for flagellar function in *P. aeruginosa.* Consistent with reports that swimming is negatively regulated by high cAMP (21), the *cpdA* mutant was swimming impaired (~53% relative to wild type). Deletion of *cpdA* in the *fimV* (~77%), *fimL* (~90%), and *pilG* (~85%) backgrounds reduced swimming relative to the single mutants, potentially due to increased cAMP levels (Figure 1). Suprisingly, despite having very high cAMP levels (17), and levels of piliation similar to the *cpdA* mutant (Figure 1), *pilH* had ~93% swimming motility relative to wild type (Figure 5). This finding suggests that high levels of cAMP and piliation do not necessarily inhibit swimming motility.

**Figure 5.**
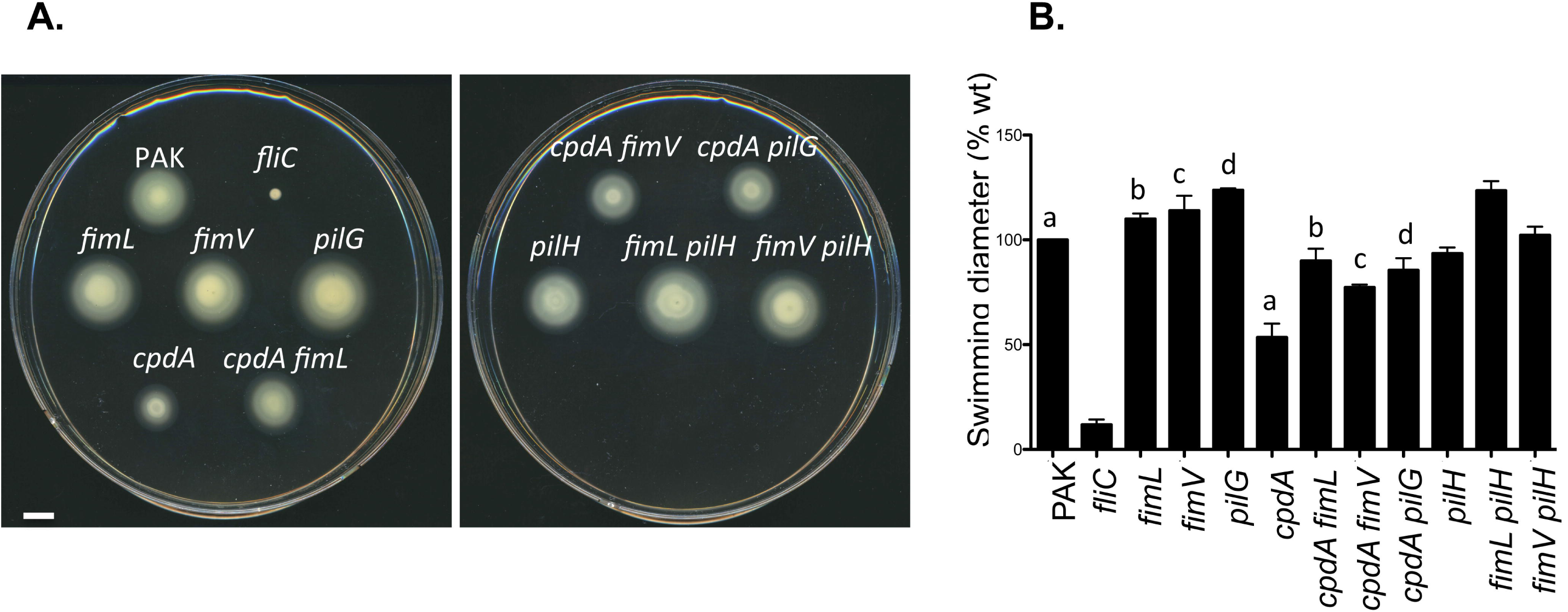
Swimming motility of PAK wild type and mutant strains. **A.** Representative swimming motility assays in 0.3% LB agar. Scale bar = 1 cm. **B.** Quantification of swimming zones, average of n = 3. Lowercase letters indicate paired samples that are statistically different from one another. Loss of *fimV* does not impact swimming motility. PAK is the parent strain; the *fliC* mutant that lacks flagellin was included for comparison.

## DISCUSSION

PilG, FimL, and FimV were recently proposed to be components of a surface-sensing pathway that activates CyaB (22), with FimL acting as a scaffold protein connecting PilG to FimV. Consistent with this model, we saw that increasing the level of PilG phosphorylation through deletion of *pilH* (18) failed to increase levels of PilU if either FimV or FimL was missing (Figure 1). However, only *the fimL* mutant twitched (Figure 1) following introduction of compensatory mutations that increased cAMP levels (17, 32), confirming that both PilG and FimV have cAMP-independent roles in twitching (17, 25). Thus, FimL's role in twitching is limited to its ability to connect PilG and FimV, leading to CyaB activation via an as-yet unknown mechanism. Interestingly, Nolan et al. (33) identified other suppressors of *fimL* that mapped outside the *cyaA*, *cyaB*, *pilG*, *pilH*, *vfr*, and *cpdA* loci. How those uncharacterized loci fit into the FimV-FimL-PilG signalling axis remains to be determined.

Since FimL's role is limited to the cAMP-dependent pathway (22, 32) and twitching motility in the *fimL* background is essentially wild type (Figure 1), PilG and FimV both function – together or independently of one another – in its absence. PilG polar localization is dependent on FimV, but PilG remains localized to the poles when FimL is absent (22). These data imply that PilG interacts with FimV directly, or indirectly via another, as-yet unidentified adaptor protein. That component is unlikely to be part of the Chp system; the MCP PilJ localizes to the poles independently of FimV, and PilG retains bipolar localization in the absence of both PilJ **(Figure S1)** and the Chp system kinase, ChpA (22). Identifying this interaction partner – potentially among the list of proteins recovered in a recent PilG pulldown/mass spectrometry study (22) could help to clarify how PilG contributes to cAMP-independent regulation of twitching.

Because restoration of cAMP levels in a *pilG* mutant by supplying exogenenous cAMP (17), constitutively activating CyaB (35), or deleting *cpdA* (Figure 1A) restores piliation but not twitching motility, the cAMP-independent role of PilG may be the coordination of pilus retraction to permit directional movement. In *M. xanthus*, the Chp-like Frz system controls the asymmetric subcellular distribution of the PilB and PilT motor ATPases in cells undergoing T4aP-mediated S-motility, to coordinate movement (39). It is likely that asymmetric T4aP retraction similarly occurs in *P. aeruginosa*, as pilus retraction at both poles simultaneously would result in zero net movement.

How PilG might regulate pilus retraction remains unclear. CheY interacts with FliM at the *E. coli* flagellar switch complex to control the direction of flagellum rotation (40), but the T4Ap system lacks an obvious FliM equivalent. However, the T4aP system was recently discovered to have a rotary motor (41–43). A hexameric PilB or PilT ATPase docks into the PilM ring at the base of the apparatus and encircles the cytoplasmic domains of the PilC platform protein (7), rotating it clockwise or counterclockwise, respectively, to insert or extract pilin subunits from the pilus in a stepwise manner (43). Transient interactions of phospho-PilG with PilM, PilC, or the motor ATPases might dictate which ATPase is docked at the leading versus lagging pole. In *pilH* mutants, hyperactivation of PilG may dysregulate asymmetric pilus retraction, leading to hyperpiliation and impaired motility (Figure 1A).

Interestingly, the cAMP-independent role of PilG appears dependent on, but distinct from, that of FimV. In addition to FimL and PilG (22), FimV is required for polar localization of the structural components PilMNOPQ (30) and the PilSR two-component system (Figure 4). However, unlike its homologues in *V. cholerae* and *S. putrefaciens* (27, 28), its deletion does not affect swimming (Figure 5). FimV and its homologs are emerging as protein interaction hubs that bind to septal PG via their LysM motif to target their partners to the septum during division, ensuring the correct placement of polar and partitioning systems during and after separation of daughter cells (28, 30). Although studies of *L pneumophila* FimV and *N. meningitidis* TspA (9, 11) did not address the role of the LysM motif or localization in function, the phenotypes of mutants lacking these proteins could reflect consequent mislocalization of motility or adhesion systems.

Septal PG binding by FimV was dispensable for CyaB activation (Figure 2) even though FimL, PilG, and CyaB are located at the cell poles (22, 32). It is possible that deletion of the LysM motif alone does not completely mislocalize FimV, as it likely has other interaction partners that help to confine it to the cell poles; attempts to test this hypothesis using a FimV_∆LysM_-YFP fusion have not been successful to date. Alternatively, FimV_∆LysM_ may be present at the cell pole due to diffusion in the inner membrane in sufficient quantities to promote CyaB activity. Consistent with only partial mislocalization of FimV_∆LysM_, PilS-YFP remained mostly localized to the poles in that background, while deletion of FimV’s entire periplasmic region led to PilS delocalization (Figure 4). Transmembrane domains 5 and 6 of PilS (44), and the membrane-embedded MASE2 domain of CyaB (35) are sufficient for their polar localization. It is possible that they interact with FimV via its transmembrane segment, or are integrated into the FimV hub through as-yet unidentified intermediaries.

In summary, this work helps to resolve the cAMP-dependent and independent regulation of *P. aeruginosa* twitching motility by FimV and PilG. The cAMP-independent role of FimV is likely coordinate localization of multiple T4aP structural and regulatory components to the cell poles, while that of PilG may be to control pilus retraction in a way that allows for directional movement; experiments to test this idea are underway. Characterization of the FimV protein-interaction network will identify its full repertoire of direct and indirect interaction partners, and clarify the links between polar localization and function.

## MATERIALS AND METHODS

### Bacterial growth and culture conditions

Bacterial strains and plasmids are listed in Table 1. Unless otherwise stated, untransformed *P. aeruginosa* strains and all *E. coli* strains were grown on LB agar at 37 °C. Antibiotic selection was as follows unless stated otherwise: gentamicin, 15μg/ml for *E. coli* and 30μg/ml for *Ρ. aeruginosa;* kanamycin, 50μg/ml for *E. coli;* ampicillin, 100mg/ml for *E. coli.* All *P. aeruginosa* strains containing a FimV complementation construct were grown on media supplemented with 0.1% (w/v) arabinose.

**Table 1.** Strains and plasmids used in this study.

| Strains | Genotype/Description | Source |
| --- | --- | --- |
| *P. aeruginosa* |  |  |
| PAK | Wild type *P. aeruginosa* strain K | (17) |
| NP | PAK with a deletion of *pilA* | (49) |
| *pilU* | PAK with a deletion of *pilU* | (25) |
| *fimV* | PAK with a deletion of *fimV* | (25) |
| *fimV*<sub>LysM</sub> | PAK with an in-frame deletion of the LysM motif, nucleotides 519-690 | This study |
| *fimV cyaB-R456L* | PAK with a deletion of *fimV* and an arginine to lysine substitution in *cyaB* at position 456 | (25) |
| *fimV1194* | PAK with an FRT insertion at nucleotide position 1194 in *fimV* | (25) |
| *fimL* | PAK with a deletion of *fimL* | This study |
| *pilG* | PAK with a deletion of *pilG* | (17) |
| *cpdA* | PAK with a deletion of *cpdA* | This study |
| *cpdA fimV* | PAK with deletions of *cpdA* and *fimV* | This study |
| *cpdA fimL* | PAK with deletions of *cpdA* and *fimL* | This study |
| *cpdA pilG* | PAK with deletions of *cpdA* and *pilG* | This study |
| *pilH* | PAK with a deletion of *pilH* | This study |
| *pilH fimV* | PAK with deletions of *pilH* and *fimV* | This study |
| *pilH fimL* | PAK with deletions of *pilH* and *fimL* | This study |
| *E. coli* |  |  |
| DH5a | F-, $\Phi 80$lacZΔM15, $\Delta$(lacZYA-argF),<br>U169, *recA1*, *endA1*, *hsdR17*(rk-,<br>*mk+*), *phoA*, *supE44*, *thi-1*, *gyrA96*, *relA1*, λ-;<br>General cloning strain | Invitrogen |
| SM10 | *thi-1 thr leu tonA lacy supE recARP42`-2 Tcr::Mu*<br>Kmr; mating donor strain | (50) |
| Plasmids |  |  |
| pBADGr | Arabinose-inducible expression construct | (51) |
| pBADGr::*fimV* | Arabinose-inducible expression construct<br>encoding full length FimV | (25) |
| pBADGr::*fimV507* | Arabinose-inducible expression construct<br>encoding the periplasmic domain of FimV,<br>residues 1-507 | This study |
| pEX18Gm::*fimL* | Suicide vector containing 1kb upstream and<br>downstream from the *fimL* locus | This study |
| pEX18Gm::*cpdA* | Suicide vector containing 1kb upstream and<br>downstream from the *cpdA* locus | This study |
| pEX18Gm::*pilH* | Suicide vector containing 1kb upstream and<br>downstream from the *pilH* locus | This study |
| pEX18Ap-*fimV*-GmFRT | Suicide vector containing *fimV* amplified from PAO1 and disrupted at nucleotide position 1194 with an FRT-flanked gentamicin resistance cassette | (29) |
| pEX18Gm-*fimV*-ΔLysM | Suicide vector containing residues 1-1521 of *fimV* with a deletion of nucleotides 519-690 | (29) |
| pBADGr::eYFP(HinDIII) | Arabinose-inducible expression construct encoding eYFP cloned into the HinDIII site | (30) |
| pBADGr::PilS-YFP | Arabinose-inducible expression construct with pilS cloned upstream of eYFP | This study |
| pMarkiC | puCP20Gm-based vector with (Gly<sub>3</sub>-Ser)<sub>3</sub>-YFP cloned in at the HinDIII site | This study |
| pMarkiC::pilG | pMarkiC with PilG cloned into the EcoRI and XbaI sites | This study |

### Mutant generation

Mutants were made as previously described (17). Deletion constructs for the generation of *fimL*, *cpdA*, and *fimV* mutants were designed to include 100 nucleotides upstream and downstream of the gene to be deleted. The *pilH* construct was designed to include the first 12 nucleotides and last 30 nucleotides of the gene. The upstream and downstream fragments were amplified from the PAK chromosome using the primer sets described in Table 2. Inserts were cloned into the pEX18Gm suicide vector at the Sacl and HinDIII sites (pEX18Gm::*fimL*), Kpnl and EcoRI sites (pEX18Gm::*cpdA*), and HinDIII and Kpnl sites (pEX18Gm::*pilH*). Suicide vectors were verified by DNA sequencing.

**Table 2.**
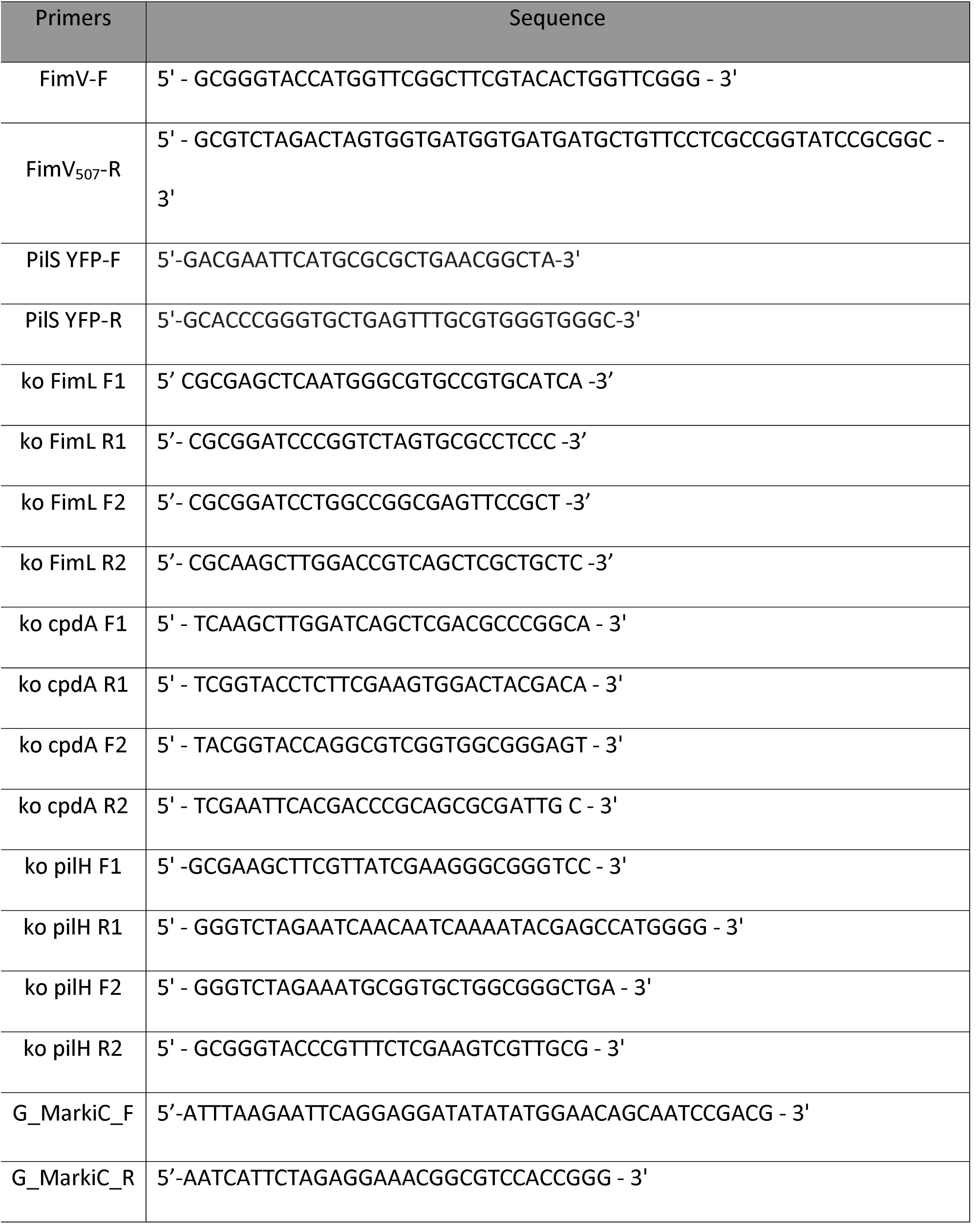
List of oligonucleotides used in this study

After verification, plasmids (pEX18GM::*fimL*, pEX18Gm::*cpdA*, pEX18Gm::*pilH*, and pEX18Gm::*fimV*_LysM_) were transformed into *E. coli* SM10 cells. Plasmids were transferred to *P. aeruginosa* by conjugation at a ratio of 6:1 (*E. coli* to *P. aeruginosa*). 100 μl of the 6:1 mixed culture were spotted onto LB 1.5% agar (w/v) and incubated overnight at 37°C. The mating mixture was resuspended in 5 ml of LB and 100 μl was plated onto *Pseudomonas* isolation agar supplemented with Gm 100 μg/ml and grown overnight at 37°C. Single colonies were resuspended in 1 ml LB and plated onto LB 1.5% agar lacking sodium chloride and supplemented with 8% (w/v) sucrose, and grown overnight at 30°C. Resulting single colonies were replica plated onto LB and LB supplemented with Gm 30 μg/ml. Mutants were verified by PCR, and the *pilH* mutant was screened by western blotting with anti-PilH antiserum. Double mutants were generated in the same manner.

### Plasmid construction

The coding region of the first 507 residues of FimV was amplified by PCR using pBADGr::*fimV* as a template, and cloned into pBADGr. The PCR amplified DNA was digested with, purified, and ligated into pBADGr at the Kpnl and Xbal sites with T4 DNA ligase according to manufacturer’s instructions (Thermo Scientific).

A version *of piIS* lacking its stop codon was amplified from the PAK chromosome and cloned into pBADGr::*yfp* in-frame with *yfp* at the EcoRI and Smal sites. In-frame ligation was confirmed by DNA sequencing.

A version of PilG lacking its stop codon was amplified from the PAO1 chromosome and cloned into the pMarkiC vector, which encodes (Gly_3_-Ser)_3_-YFP at the HinDIII site of pUCP20Gm. PilG was ligated into the EcoRI and Xbal sites. In-frame ligation was confirmed by DNA sequencing.

### Immunoblotting

Western blotting of whole cell lysates was performed as previously described (45). In brief, whole cell lysates were prepared from strains were grown overnight on LB 1.5% agar, or in the case of plasmid transformed strains, LB 1.5% agar supplemented with 0.1% (w/v) arabinose. Cell growth was then resuspended in 1X PBS and normalized to an OD_600_ of 0.6. Cells were pelleted by centrifugation at 2,300 ×*g* for 5 min. Pellets were then resuspended in 175 μl of 1X SDS-PAGE loading dye. Cell lysates were resolved on 15% SDS-PAGE gels and transferred to nitrocellulose membranes. Membranes were blocked in 5% skim milk dissolved in PBS (pH 7.4) for 1 h, washed in PBS, and incubated with PBS-diluted antisera raised against the FimV periplasmic domain (1:1000), PilU (1:5000), PilM (1:1000), PilN (1:1000), PilO (1:1000), PilP (1:1000), or PilQ (1:1000), or polyclonal anti-GFP antibody (Novus Biologicals; 1:5000) for 1 h, washed, incubated with alkaline phosphatase-conjugated goat-anti-rabbit secondary antibody (1:3000, Bio-Rad) for 1 h, and washed. Blots were developed using 5-bromo-4-chloro-3-indolylphosphate (BCIP) and nitro blue tetrazolium (NBT). Data are representative of n = 3 independent experiments.

### Sheared surface protein preparation

Surface pili were analyzed as previously described (46). In brief, strains of interest were streaked in a grid-like pattern onto LB 1.5% agar, or in the case of plasmid-transformed strains, LB 1.5% agar supplemented with 0.1% (w/v) arabinose and grown overnight at 37 °C. Cells were gently scraped from the plates using a sterile coverslip and resuspended in 4.5 ml PBS (pH 7.4). Surface appendages were sheared by vortexing the cells for 30 s. The OD_600_ for each strain was measured, and an amount of cells equivalent to 4.5 ml of the sample with the lowest OD_600_ was pelleted by centrifugation at 16,100 × *g* for 5 min. When necessary, PBS was added to samples to a final volume of 4.5 ml prior to centrifugation. Supernatants were removed and centrifuged again at 16,100 × *g* for 20 min to remove remaining cells. Supernatants were collected and mixed with 5 *M* NaCl and 30% (w/v) polyethylene glycol (Sigma; molecular weight range ~8000) to a final concentration of 0.5 *M* NaCl and 3% (w/v) polyethylene glycol, and incubated on ice for 30 min. Precipitated surface proteins were collected by centrifugation at 16,100 x g for 30 min. Supernatants were discarded and samples were centrifuged again at 16,100 x g for 2 min. Pellets were resuspended in 150 μl of 1X SDS-PAGE sample buffer (80 m*M* Tris, pH 6.8, 5.3% (v/v) 2-mercaptoethanol, 10% (v/v) glycerol, 0.02% (w/v) bromophenol blue, 2% (w/v) SDS). Samples were boiled for 10 min and resolved by 15% SDS-PAGE. Bands were visualized by staining with Coomassie brilliant blue (Sigma). Data are representative of n = 3 independent experiments.

### Twitching assay

Twitching motility was tested as previously described (46). In brief, cells from an overnight culture were stab inoculated to the interface between LB 1% agar, or in the case of plasmid-transformed strains, LB 1% agar supplemented with 0.1% (w/v) arabinose and the underlying tissue culture-treated polystyrene petri dish, and incubated at 37 °C for 16 h (Thermo Fisher). Twitching zones were visualized by removing the agar and staining cells on the petri dish with 1% (w/v) crystal violet and washing with water to remove unbound dye. Twitching zones were measured by analyzing the diameter of each twitching zone in pixels using ImageJ software (NIH). Twitching zones were normalized to the twitching diameter of wild type PAK in each individual experiment. Data are representative of n = 3 independent experiments.

### Fluorescence microscopy

*P. aeruginosa* strains transformed with pBADGr::FimV-eYFP were grown overnight. Microscopy was performed using 8-well 1.0 borosilicate chambered coverglass (LabTek). Chamber slides were prepared by adding LB 1% agar supplemented with 0.1% (w/v) arabinose to create an agar layer ~3mm in thickness and covering the bottom of the chamber. Agar was allowed to solidify with the lid off to prevent condensation. Bacteria were stab inoculated to the interface between the agar and coverglass. Slides were wrapped in foil to prevent photobleaching, and incubated at 37°C for 1h in the dark. Cells were then imaged using an EVOS FL Auto microscope, with a monochrome camera for brightfield imaging and a YFP LED light cube for fluorescence imaging, through a 60X oil immersion objective at room temperature. Representative fields were cropped from larger images and enlarged using ImageJ software (NIH) (47).

Fluorescence images were quantified using the MicrobeJ plugin for ImageJ (48). Brightfield and fluorescence images were arranged into a stack on ImageJ. Regions of interest corresponding to the bacteria were selected based on the brightfield image, and thresholding particles based on length (0.5μm-5μm), width (0.2μm-1.5μm), and area (0.75μm^2^-max), and fit to rod-shaped bacteria. Pixel intensity profiles were generated by MicrobeJ using the profile option on the fluorescence image, using 1μm width and 0.5μm extensions. Intensity profiles were plotted along a Y-axis of range 0-140, and the X-axis was partitioned into 50 bins. Pixel intensity profiles were generated for the YFP channel. Data are representative of at least 3 independent trials.

### Swimming assay

Cells from overnight cultures were resuspended in sterile PBS and standardized to OD_600_ 0.6. Two μl of cell suspension were spotted onto LB 0.3% agar and allowed to dry onto the surface of the agar. Plates were incubated at 30°C for 16h with the agar side down. Data are representative of n=3 independent experiments.

